# RasGRP1 Signalling Programs Developing γδ-Thymocytes towards the γδT17 Lineage Through Control of c-Maf Expression

**DOI:** 10.1101/2024.12.13.628252

**Authors:** K. Joannou, D.P. Golec, L.M. Henao-Caviedes, J.F. May, R.G. Kelly, T.A. Baldwin

## Abstract

The γδ TCR instructively directs both lineage specification and effector programming of developing γδ T cells. However, the manner in which different TCR signal strengths and other auxiliary signals coordinate downstream of the γδ TCR to regulate γδ T cell development remains unclear. In this study we characterized the role of Ras guanyl-releasing protein 1 (RasGRP1) in the development and effector programming of γδ T cells. While RasGRP1 was not necessary for bulk γδ T cell generation, we found it required for efficient generation of Vγ4^+^ thymocytes, and lineage-committed CD73^+^ γδ T cells in the thymus and periphery. Despite a decrease in immature CD73^+^ γδ thymocytes, we report an expansion of the perinatal wave of CD8^+^IFNγ^+^ γδ T cell population in the absence of RasGRP1. IL-17 producing γδ T cells were significantly reduced in RasGRP1 KO mice, with a specific loss of Vγ2^+^ γδ T cells that corresponds to a loss of c-Maf expression as early as the DN1d thymocyte stage. Critically, these adult-programmed γδT17s could express c-Maf in response to CCR9 stimulation, with RasGRP1 being required for CCR9-induced c-Maf expression. Thus, RasGRP1 activation serves as an important signalling hub in the effector programming of γδ T cells, which integrates signals from both non-TCR and TCR inputs to direct differentiation.

## Introduction

T cell development is a complex process that starts with hematopoietic stem cells and terminates with mature functional subsets of T cells. This process includes many layers of regulation to ensure that T cells are directed into specific lineages and in properly balanced numbers based in part on their T Cell Receptor (TCR) specificity. Therefore, for many T cell subsets, TCR signals received by progenitors critically regulate development and differentiation.

αβ T cells undergo a developmental process that includes both clonal selection and lineage direction into subtypes of conventional or agonist-selected populations based on TCR interactions with self-peptide-MHC. In contrast, many γδ T cell subsets in mice have poorly defined ligands, and do not seem to require classical MHC or peptide-MHC complexes for their development^1,2^. This mystery has led to difficulty in defining how TCR signals direct γδ T cell development. Current evidence suggests that the γδ TCR delivers a stronger signal to developing thymocytes compared to the pre-TCR. This strong γδ TCR signal directs cells into the γδ lineage instead of the αβ lineage^3–5^. In addition, stronger γδ TCR signals received during development have been shown to direct developing γδ T cells into IFNγ producing (γδT1) over IL-17 producing (γδT17) subsets^6,7^. Critically, Erk activation seems to bias development towards γδT17 programming^8^.

The roles of different signaling pathways or regulators of these pathways downstream of the γδ TCR in translating the TCR signal into distinct lineage and effector directions remain unclear. The Ras pathway is primarily activated downstream of the TCR in developing T cells through the Ras guanine nucleotide exchange factors (GEFs) RasGRP1 or Son of Sevenless 1 (SOS1)^9^. Through the trajectory of αβ T cell development, SOS1 is more highly expressed early during double-negative (DN) stages, whereas RasGRP1 expression is upregulated through the DN stages and becomes more highly expressed by the double-positive (DP) stage^10^. Both RasGRP1 and SOS1 contribute to both β-selection and negative selection, but efficient positive selection requires RasGRP1 but not SOS1^11–13^. Agonist selection is dependent on RasGRP1, but it is unclear whether SOS1 is necessary^13^. Critically, SOS1 has some role in producing normal numbers of γδ thymocytes^10^, but how specific Ras GEFs contribute to different developmental checkpoints for γδ T cells remains poorly characterized. Furthermore, it remains unknown how different Ras activators contribute to γδ effector lineage programming, but some evidence suggests RasGRP1 transmits signals that somehow bias this process towards γδT17 programming^14^.

Previous research on RasGRP1 knockout (KO) mice determined that without RasGRP1 there are more total γδ T cells in the periphery and a shift away from γδT17s and towards γδT1s^14^. However, the mechanisms to explain how RasGRP1 contributes to normal γδ T cell populations is currently unknown. In this study we investigated the roles of RasGRP1 in γδ T cell development and effector lineage programming. We found that RasGRP1 influences the proportion of Vγ4^+^ in the neonatal thymus, and proportions of Vγ1.1^+^ and Vγ2^+^ γδ T cells in the adult thymus (Garman nomenclature)^15^. We also discovered that RasGRP1 signaling constrains CD8^+^ γδT1 development during early life. Furthermore, RasGRP1 signals from non-TCR sources begin γδT17 programming in the DN1d stage before TCR expression and CCR9 stimulation can induce c-Maf expression in γδ-thymocytes in a RasGRP1 dependent manner.

## Materials and Methods

### Mice

All mice were bred and maintained in our colony at the University of Alberta and on the C57BL/6 background. Rag2pGFP^16^ mice were provided by Dr. Pamela Fink. Nur77-GFP^17^ mice were provided by Dr. Colin Anderson. Both male and female mice were analyzed between 8 and 14 wk of age for all experiments. All mice were treated in accordance with protocols approved by the University of Alberta Animal Care.

### Bone Marrow Chimeras

Donor mice were depleted of T cells by injection of 100 μg of purified 30H12 (anti-Thy1.2) Ab (Bio X Cell) on days −2 and −1. Recipient wild-type mice were lethally irradiated with 2× 450 rad separated by 1–4 h on day −1 and transplanted on day 0 with 5–10 × 10^6^ donor bone marrow cells at a mixture of a 1:4 mixture of WT:RasGRP1 KO. Mice were provided sulfamethoxazole/trimethoprim (Novo-Trimel) in their drinking water for 28 d posttransplant.

### Tissue Collection and Cell Isolation

Thymus, spleen, and lymph nodes were ground through wire mesh or nylon screens into sterile PBS or EZSep Buffer (PBS, 1% FCS, 1mM EDTA) to create single-cell suspensions. Mesenteric and inguinal lymph nodes were mixed together unless otherwise stated. To collect dermal lymphocytes, ears were harvested, separated into dorsal and ventral halves, and cut into <2-mm pieces in ice-cold PBS. Skin was then digested for 2 h at 37°C in DMEM with DNase I (5 μg/ml) and collagenase (2 mg/ml). Cells were disaggregated by pipetting every 30 min, and passed through a 70-μm filter into ice-cold RPMI 1640 medium with 0.5 mM EDTA at the end of digestion.

### Antibodies and Flow Cytometry

Fluorochrome-conjugated and biotinylated Abs were purchased from BD Biosciences, BioLegend, and ThermoFisher Scientific. Specific antibody clones used were CD24(M1.69), CD73(TY/11.8), TCRδ(GL3), CD3ε(145-2C11), Ki67(SolA15), CD45.1(A20), CD45.2(104), CD4(RM4-5), CD8α(53-6.7), TER-119(TER-119), GR1(RB6-8C5), CD19(1D3/CD19), CD11b(M1/70), CD11c(N418), IFNγ(XMG1.2), IL17A(TC11-18H10.1), TCRβ(H57-597), Vγ1.1(2.11), Vγ2(4C3-10A6), Vγ3(536), c-Maf(sym0F1), RORγt(Q31-378), CD44(IM7), CD62L(MEL-14), CCR9(eBioCW-1.2), NK1.1(PK136). Live/Dead staining was done using Ghost Dye Violet 510 (Tonbo Bioscience) or Live/Dead Fixable Near-IR Dead Cell Stain Kits (Invitrogen), and was applied to all experiments that involved *in-vitro* culture or stimulation. Cells were incubated with the Live/Dead stain for 5 minutes in PBS on ice followed by 2 washes in FACS Buffer (PBS, 1% FCS, 0.02% sodium azide, 1 mM EDTA). Ab cocktails were applied to cells in FACS buffer for 30 min on ice in the dark, and were washed twice with FACS buffer between staining steps. A pre-fixation of cells expressing GFP was done with 2% formaldehyde in PBS for 45 minutes prior to permeabilization to preserve GFP intensity. Permeabilization for intracellular antigen staining was done with a FoxP3 Transcription Factor Staining Buffer Set (ThermoFisher Scientific). Flow cytometry was performed on a BD LSRFortessa (BD Biosciences), BD LSRFortessa X-20 (BD Biosciences), or Attune NXT (Invitrogen), and data were analyzed with FlowJo v10.6.2 (BD Biosciences).

### Cytokine Induction

Bulk thymocytes or splenocytes were treated with 3 μg/ml brefeldin A (Invitrogen), without or with Phorbol 12-myristate 13-acetate (PMA) (20 ng/ml) and ionomycin (IM) (1 μg/ml), and incubated at 37°C, 5% CO_2_ for 3 or 4 h in sterile RP10 media (RPMI 1640 with 2.05 mM l-glutamine [HyClone], 10% FCS, 5 mM HEPES, 50 μM 2-ME, 50 mg/ml penicillin/streptomycin, and 50 mg/ml gentamicin sulfate) prior to preparation to intracellular cytokine staining as described above.

### Statistical Analysis

Statistical analysis was performed using R or GraphPad Prism. Mean and SD were calculated with the plyr package (v1.8.6). Error bars represent 1 SD. Dot plots were generated and two-tailed unpaired Student *t* tests were performed using the ggpubr package (v0.4.0). Statistically significant differences between groups are noted as follows: \**p* < 0.05, \*\**p* < 0.01, \*\*\**p* < 0.001, \*\*\*\**p* < 0.0001.

## Results

### RasGRP1 KO Mice have altered populations of γδ-thymocytes and mature peripheral γδ T cells

It was previously reported that RasGRP1 KO mice have changes to bulk γδ T cell numbers and a bias towards IFNγ producers ^11,14^, but the mechanism underlying these alterations was unclear. Flow cytometric analyses confirmed normal numbers of γδ thymocytes in RasGRP1 deficient mice, but an expanded γδ T cell population in the periphery (Figure 1A). An increased frequency of γδT1 and decrease in γδT17 thymocytes was also observed (Figure 1B). We hypothesized that there were broad changes to γδ T cell development in RasGRP1 KO mice, therefore, we sought to further characterize the thymic and peripheral populations by analyzing the expression of CD24 and CD73 (Figure 1C, Supplementary 1A). CD24 downregulation is associated with functional competence and maturity in γδ T cells^18,19^, and CD73 expression is associated with stronger TCR signals during development and γδT1 programming^6^. Within the thymus, we observed a significant loss of CD24^+^CD73^+^ (Immature CD73^+^) γδ T cells in RasGRP1 KO mice, which corresponded with a proportional increase in the less mature CD24^+^CD73^-^ (Immature CD73^-^) population (Figure 1D-E). In the periphery, the Immature CD73^+^ γδ T cell population was also absent in RasGRP1 KO mice, and a comparably sized increase in Immature CD73^-^ γδ T cells was present (Figure 1D, F). Furthermore, the expansion of γδ T cell numbers among splenocytes was concentrated in the mature populations. We did note that in RasGRP1 KO CD24^+^ thymocytes there was an increase in Vγ1.1^+^ and corresponding decrease in Vγ2^+^ γδ T cells, but this change was not apparent in mature γδ thymocytes or bulk γδ splenocytes (Figure G-H, Supplementary 1A). Critically, despite a known loss of γδT17s in RasGRP1 KO mice, peripheral Vγ2^+^ and Vγ1,2,3^-^ (herein called Vγ4^+^) populations appeared normal (Figure 1H). Overall, these data suggest that RasGRP1 is not necessary for bulk development of γδ thymocytes in the adult thymus, but is required for differentiation of CD73^+^ γδ thymocytes, and has influences the proportion of immature γδ thymocytes that are Vγ1.1^+^ or Vγ2^+^.

**Figure 1:**
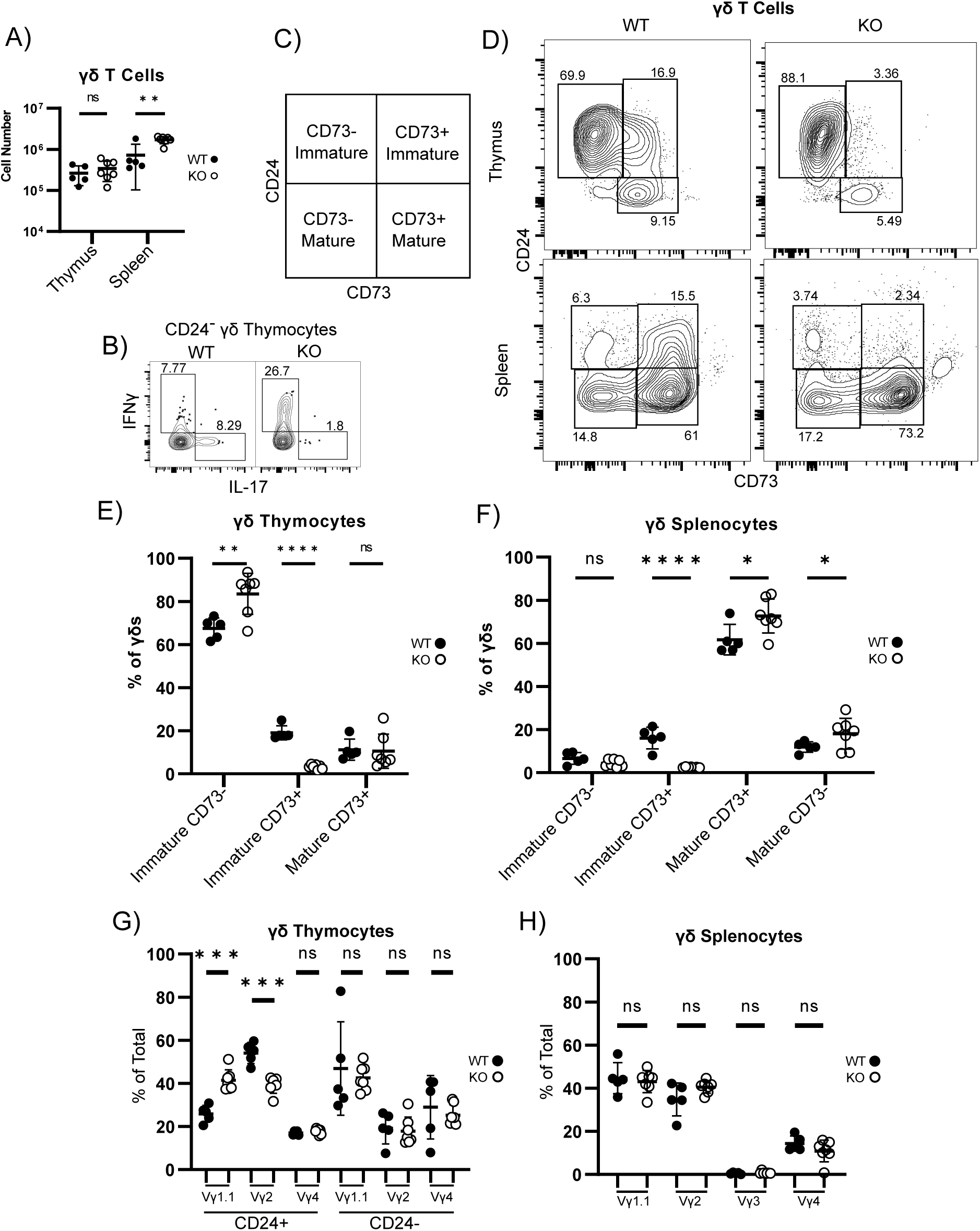
Expanded Mature γδ T cell population in RasGRP1 KO Mice. A) Total γδ T cell populations in thymus and spleen of WT and KO mice. B) Representative FACS plots of IL-17 and IFNγ production in mature γδ thymocytes after PMA/IM stimulation C) Diagram of γδ T cell maturity gating strategy. D) Representative plots of WT and KO γδ thymocytes and splenocytes. E-F) Proportion γδ T cells in different maturity stages in thymocytes (E) and splenocytes (F). G-H) Proportion of γδ thymocytes (G) or γδ splenocytes (H) that express different Vγ chains in WT or RasGRP1 KO mice. N = 5 (WT) or 7 (KO), 4 independent experiments.

### RasGRP1 KO mice have expanded mature circulating γδ T Cell population despite an intrinsic developmental impairment

We next sought to explain the expansion of bulk mature γδ T cells in the periphery of RasGRP1 KO mice. Since RasGRP1 KO mice did not appear to have a developmental advantage, we reasoned that this expansion of mature γδ T cells in the periphery could be due to a cell intrinsic or extrinsic proliferation effect. Therefore, we generated mixed competitive bone marrow chimeras (BMCs) to test for a cell intrinsic RasGRP1 KO advantage to populate the periphery. CD45.2+ RasGRP1 KO bone marrow was mixed with CD45.1+ bone marrow at a 4:1 ratio and transplanted into lethally-irradiated CD45.1/2+ recipients. A minimum of 8 weeks later, the thymus and spleen were harvested and the ratio of RasGRP1 KO to WT γδ T cells was calculated. WT advantage multiplier was calculated based on the chimerism ratio of CD44^+^cKit^+^ early thymic progenitors (ETPs) (Supplementary 1C) compared to the ratio in populations of interest. These experiments demonstrated a ∼5x advantage to WT cells populating the immature CD73^-^ thymocyte compartment, and an ∼50x advantage to CD73^+^ thymocyte populations (Figure 2A). Among splenocytes, WT cells also dominated the CD73^+^ populations, but had only a small advantage in CD73^-^ populations (Figure 2B). These data are consistent with a more minor role for RasGRP1 in CD73^-^ γδ T cell development, whereas efficient generation of CD73^+^ γδ T cell populations are highly dependent on RasGRP1 in the thymus and spleen. Given there was a cell intrinsic disadvantage to mature RasGRP1 KO γδ T cells populating the periphery, we instead hypothesized that the lymphopenic environment caused by arrested αβ T cell development in RasGRP1 KO mice causes the γδ T cell population to expand. Consistent with this hypothesis, we detected elevated Ki-67 expression in the mature, but not immature population of mature circulating γδ T cells in intact RasGRP1 KO mice (Figure 2C).

**Figure 2:**
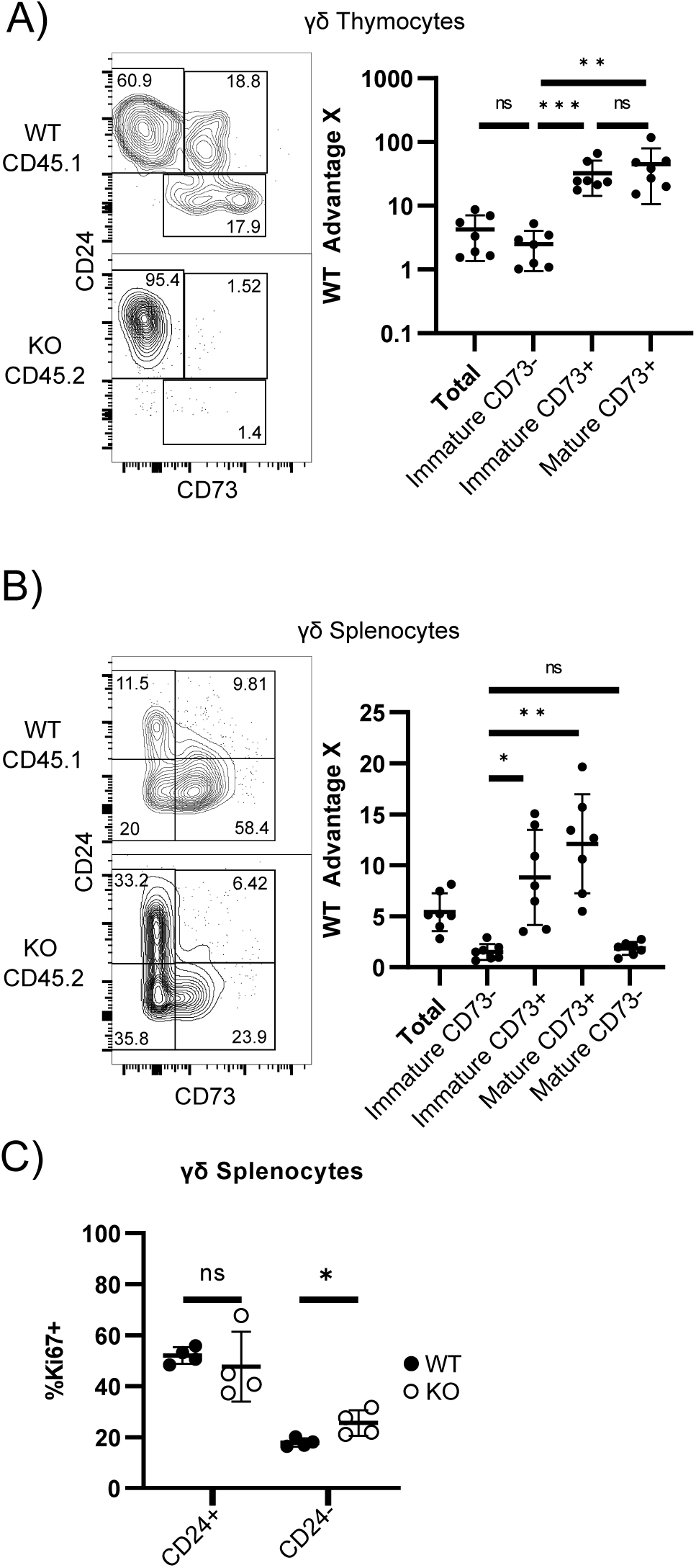
Increase in mature γδ T cells in RasGRP1 KO mice not due to improved generation. A-B) Ratio of WT/KO γδ T cells repopulating competitive mixed BMC mice. Representative FACS plots (left) and dot plots (right) of ratio of WT/KO ETPs compared to ratio in maturing populations of A) thymocytes or B) splenocytes to find multiple of WT advantage. N=7, 6 independent experiments from 3 cohorts of BMCs. C) % of mature γδ splenocytes that are Ki67+ in Immature and mature populations in WT vs RasGRP1 KO mice. N=4, 4 independent experiments.

### RasGRP1 KO mice can generate naive phenotype γδ T cells and maintain T_EM_ phenotype populations

Since RasGRP1 KO mice have altered γδ effector populations, we hypothesized that RasGRP1 KO mice may also have changes in the activation of states of circulating γδ T cells. Analysis of γδ splenocytes revealed similar proportions, yet increased numbers, of naive CD44^-^Vγ2^+^ or Vγ1.1^+^ γδ T cells in RasGRP1 KO spleens relative to WT counterparts (Figure 3A-C). In contrast, the populations of Vγ2^+^ and Vγ1.1^+^ γδT_CM_ (CD44^+^CD62L^+^) were proportionally and numerically increased in RasGRP1 KO mice, while γδT_EM_ (CD44^+^CD62L^-^) populations were proportionally lost in RasGRP1-deficient mice despite maintaining WT cellularity (Figure 3A-C). Furthermore, when we analyzed γδ dermatocytes as a representative tissue-resident population, we found that the Vγ2^+^ population of γδ T cells was absent (Figure 3D).

**Figure 3:**
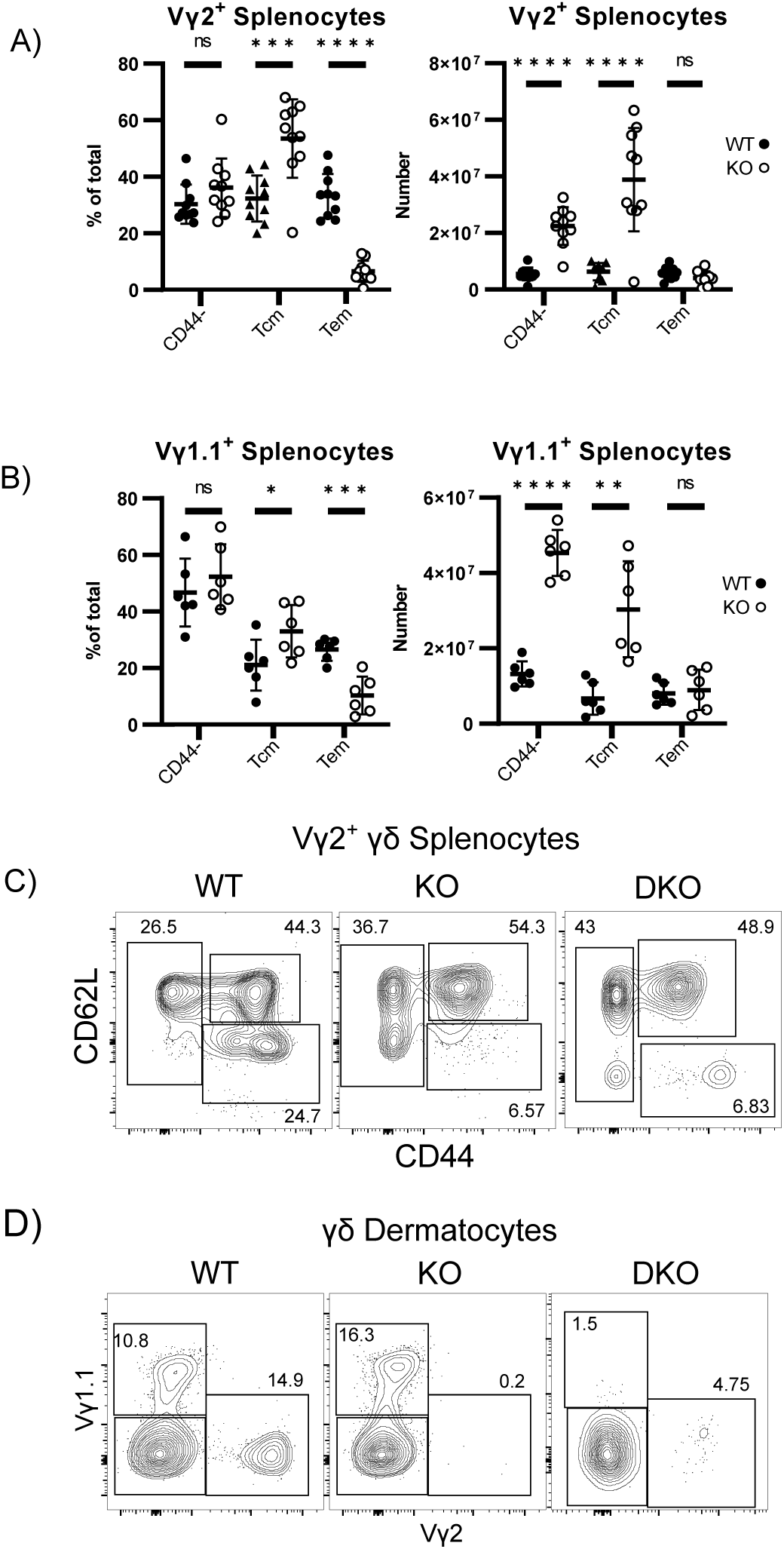
Alterations in the composition of circulating and dermal γδ T cells in RasGRP1 KO mice. A-B) Proportion and total number of Vγ2+ (A) or Vγ1.1+ (B) γδ splenocytes that are Naive, Tem, or Tcm in phenotype. N = 10 (Vγ2) or 6 (Vγ1.1), 5 or more independent experiments. C) FACS plots of CD44 and CD62L expression on WT, RasGRP1 KO, and RasGRP1/BIM DKO γδ splenocytes. D) FACS plots of proportion of Vγ1.1+ and Vγ2+ in γδ dermatocytes. C-D representative of 3 independent experiments.

Since RasGRP1 KO mice could produce naive γδ T cells but showed reduced Vγ2^+^ γδTem cells in spleen and dermis, we hypothesized that the T_EM_ population may be missing due to a requirement of RasGRP1 for survival. To test this hypothesis we analyzed RasGRP1/Bim double-knockout (DKO) mice to determine if the circulating or dermal γδT_EM_ population was rescued when Bim dependent apoptosis was removed. The γδT_EM_ population was not rescued in DKO mice (Figure 3C-D), refuting the hypothesis that a failure in homeostatic maintenance explains reduced γδT_EM_ populations in RasGRP1 KO mice.

### Expanded γδT1 population in RasGRP1 KO mice is a neonatally derived CD8^+^ population

We and others found an expansion of γδT1s in RasGRP1 KO mice (Figure 1B and^14^), but the mechanism surrounding the generation of this population was unclear. Analysis of spleen and LN showed an increase of γδT1s specifically among γδ lymphocytes, but not γδ splenocytes in RasGRP1 KO mice, which we further determined to be an expansion of the recently discovered CD8^+^ population of γδT1s^20^(Figure 4 A-B). Since these CD8^+^ γδT1 cells were shown to be generated early in life, we analyzed mixed BMCs for this CD8^+^ γδT1 population. While the CD8^-^ γδT1 population was regenerated from RasGRP1-deficient progenitors, the RasGRP1-deficient CD8^+^ γδT1 population did not substantially regenerate (Figure 4B-C). These data suggest RasGRP1-dependent ERK activation is inhibitory for the generation of CD8^+^ γδT1 cells.

**Figure 4:**
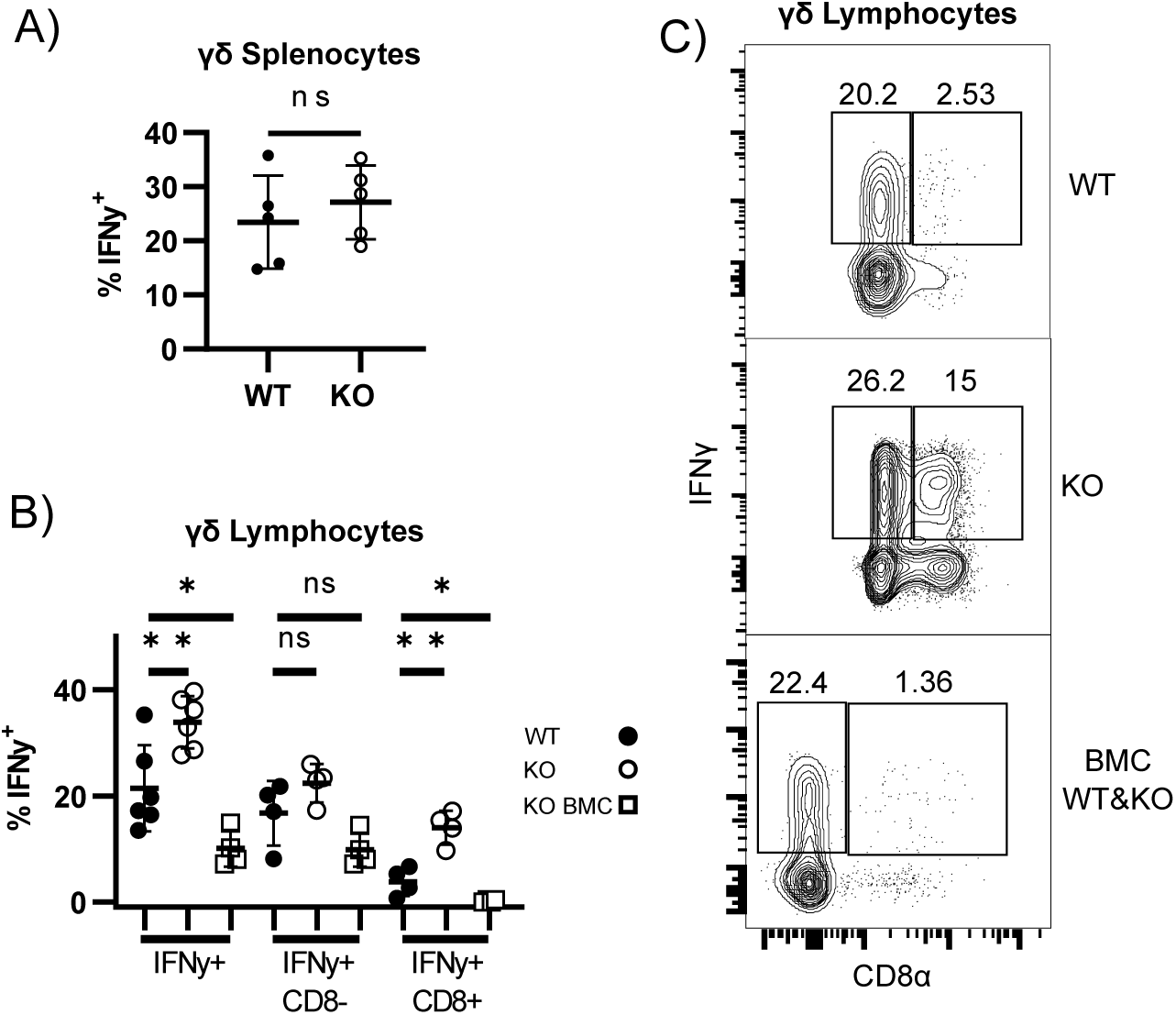
Increase in γδT1s in RasGRP1 KO mice are an expansion of neonatally-derived CD8+ cells. A-C) % of γδ Splenocytes or B-C) γδ Lymphocytes (inguinal and mesenteric mixed) producing IFNγ following 3 hours of PMA/IM stimulation. B-C) Dot and representative FACS plots of IFNγ producing γδ lymphocytes from WT, RasGRP1 KO, and the KO cells from WT/KO -> WT mixed BMCs. All Experiments N = 4 or more and 3 or more independent experiments.

### RasGRP1 regulates the generation of Vγ2^+^ γδT17 populations in the thymus and periphery

RasGRP1 KO mice have a substantial loss of circulating γδT17s^14^. It is thought that many γδT17s are produced during the fetal and neonatal periods in successive developmental waves of primarily Vγ4^+^ and Vγ2^+^ subsets^21^. Therefore, we sought to determine if Vγ4^+^ and Vγ2^+^ γδT17s may be differentially affected by RasGRP1 deficiency. A wide analysis of circulating and tissue-resident γδ T cell populations stimulated with PMA/IM demonstrated a significant loss of γδT17s in all tissues analyzed by proportion of IL-17^+^ γδ T cells (Figure 5A). However, organ specific quantitation of γδT17 numbers did not reveal a significant difference (Figure 5B). Analysis of Vγ4^+^ and Vγ2^+^ populations of γδT17s revealed that the γδT17 cells in RasGRP1 KO mice were primarily Vγ4^+^, whereas Vγ2^+^ γδT17 populations were substantially reduced (Figure 5C, E). Interestingly, quantitation of Vγ4^+^ γδT17 splenocytes appears to show a replacement of the missing Vγ2^+^ γδT17 cells with Vγ4^+^ γδT17 cells (Figure 5D-F). Taken together, these data demonstrate that Vγ4^+^ γδT17s are far less dependent on RasGRP1 for their presence in tissues or circulation than Vγ2^+^ γδT17s.

**Figure 5:**
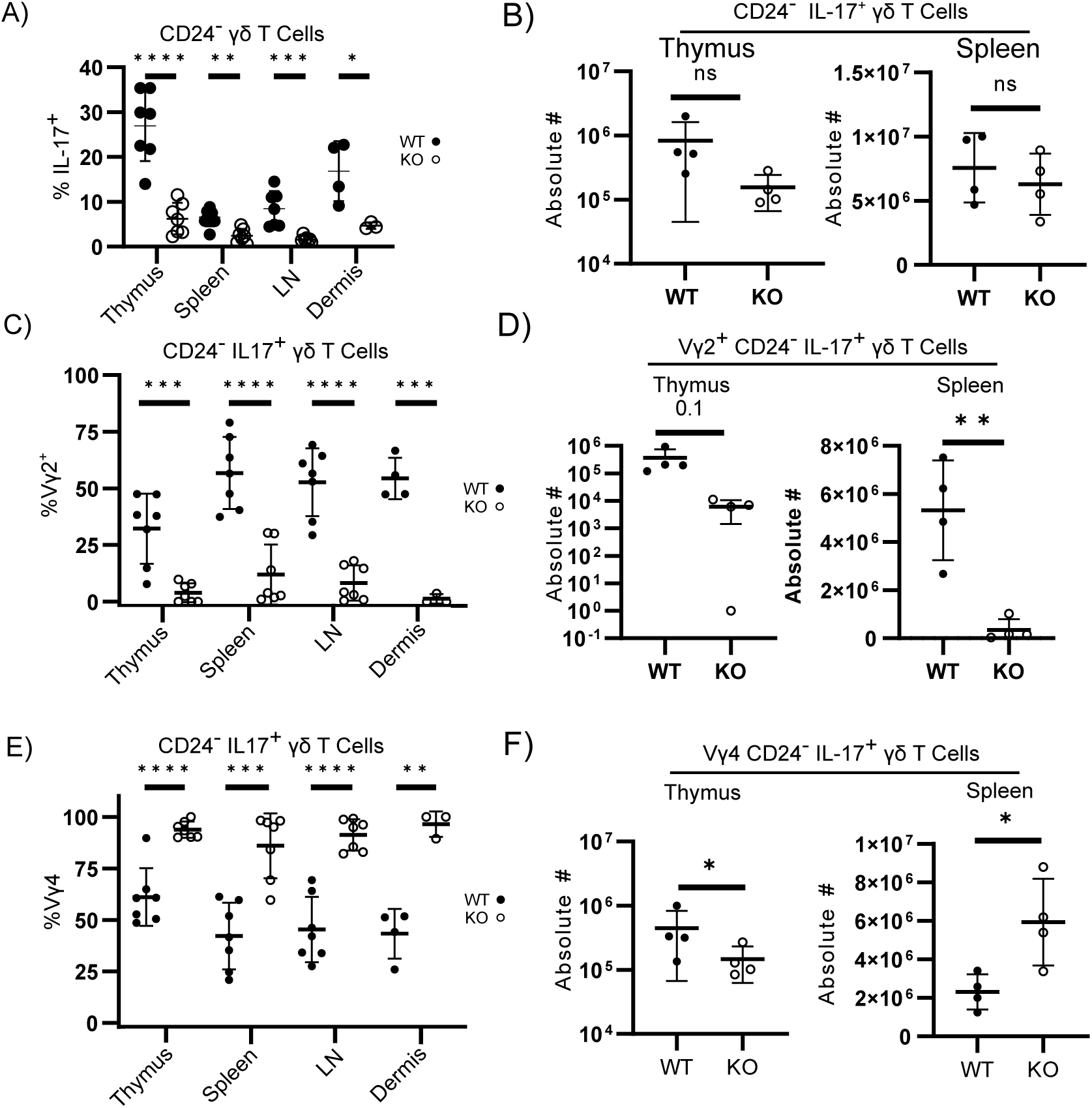
RasGRP1 KO mice lack Vγ2+ γδT17s. A-F) Percent (A, C, E) and absolute number (B, D, F) of CD24- γδ T cells in different organs that are (A-B) IL17+, (C-D) IL17+ Vγ2+ or (E-F) IL17+Vγ4+. N = 4 or more, 2 or more independent experiments.

### Vγ2^+^ γδT17 and bulk Vγ4^+^ development is perturbed in the neonatal period

The neonatal period is an important time for the development of γδT17 cells. Therefore, we extended our analysis of γδT17 differentiation in RasGRP1-deficient thymi to neonatal day 1-2. Quantitation of total γδ thymocytes in neonatal thymi demonstrated no significant change to bulk γδ or Vγ2^+^ thymocyte numbers, however, Vγ4^+^ numbers were reduced by about half (Figure 6A). Analysis of broad developmental states marked by CD24 and CD73 expression demonstrated the same loss of immature CD73^+^ γδ thymocytes as seen in adult mice (Figure 6B). There was also an increase in immature CD73^-^ γδ thymocytes in RasGRP1 KO thymi that corresponds with a decrease in mature CD73^-^ “Pathway 2”^22^ γδ thymocytes, but surprisingly, we found no change in mature CD73^+^ cells (Figure 6B). Cytokine analysis revealed a drastic decrease in γδT17 cells in RasGRP1 KO neonatal thymi, with the majority of these cells being Vγ4^+^ (Figure 6C-D). Interestingly, the proportion of γδT17 cells that were Vγ4^+^ did not change in the RasGRP1 KO, but there were so few remaining Vγ2^+^ γδT17 cells that the minor Vγ1.1^+^ γδT17 population showed a higher frequency than the Vγ2^+^ γδT17 population (Figure 6D). Analysis of the frequency of IL-17^+^ cells within each Vγ population revealed that Vγ2^+^ γδT17 cells were most heavily affected by RasGRP1 deficiency, but all populations generated γδT17s less efficiently (Figure 6E). Together these data suggest that neonatal γδT17 development is perturbed by loss of RasGRP1, such that Vγ2^+^ cells are seldom programmed to be γδT17s, while Vγ4^+^ cells are capable of being programmed into γδT17s, albeit with reduced efficiency.

**Figure 6:**
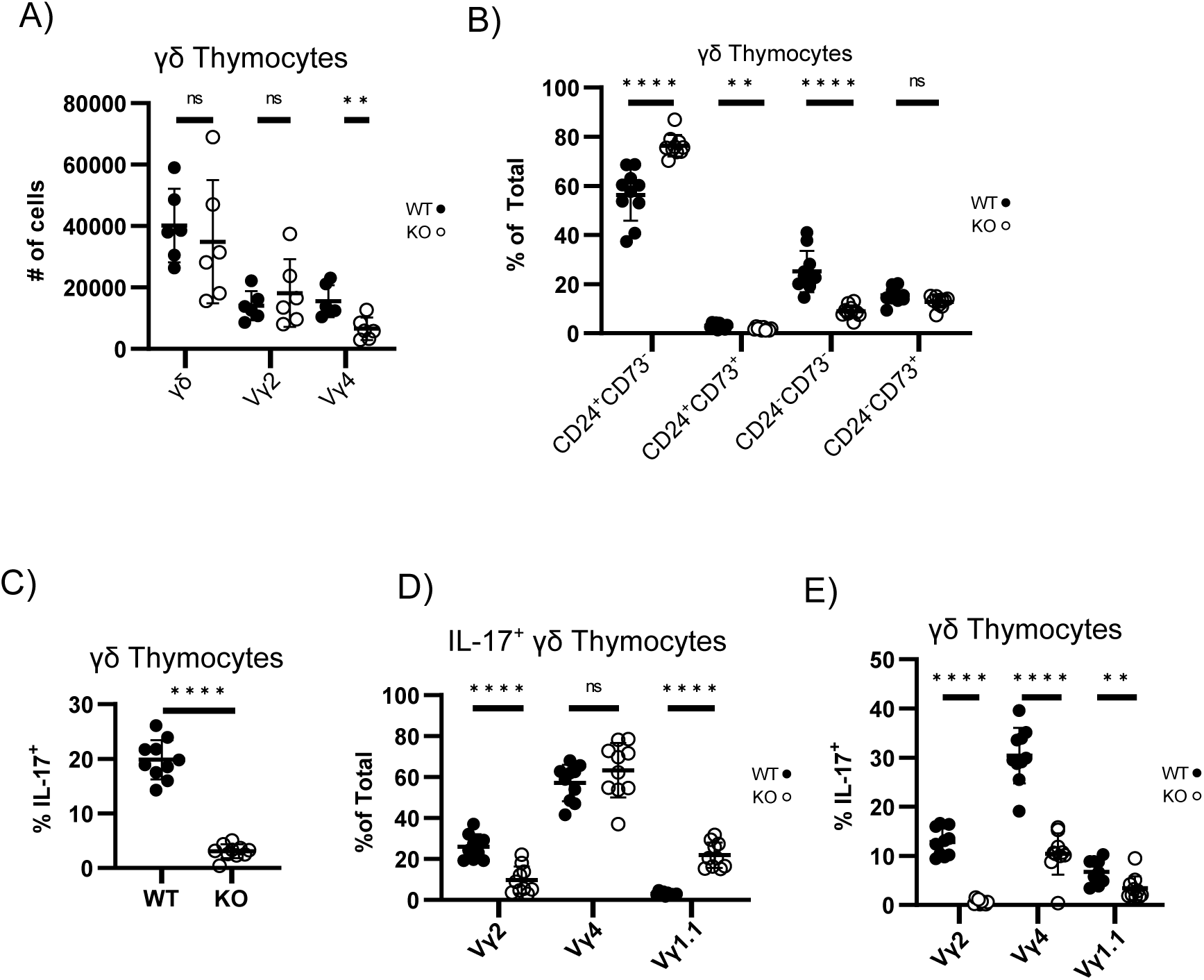
Vy2+ γδT17 development is perturbed in the neonatal period. A) Bulk, Vγ2+, and Vγ4+ thymus cellularity in D1-2 WT and RasGRP1 KO mice. B) Proportion of γδ thymocytes in CD24/CD73 quadrants in D1-2 WT and RasGRP1 KO mice. C) Proportion of γδ thymocytes producing IL-17 in D1-2 WT and RasGRP1 KO Mice after 3 hours PMA/IM stimulation. D-E) Proportion of D) IL-17+ γδ thymocytes that are Vγ2+, Vγ4+, or Vγ1.1+, or E) % of Vγ2+, Vγ4+, or Vγ1.1+ γδ thymocytes that are IL-17+ in D1-2 WT and RasGRP1 KO mice after 3 hours PMA/ IM stimulation. N=6 or 10, 2 or more independent experiments.

### γδT17 programming defect is detectable among immature Vγ2^+^ γδ T cells

We next sought to determine if we could identify a specific developmental stage in Vγ2^+^ γδT17 programming that is blocked in the absence of RasGRP1. Therefore, we analyzed immature Vγ2^+^ γδ T cells in the thymus and periphery for expression of the master transcription factors (TFs) that regulate IL-17 production: c-Maf and RORγt. We found a clear c-Maf^+^RORγt^+^Vγ2^+^ thymocyte population, which even at the CD24^+^ stage showed a significant loss of both c-Maf and RORγt expression in RasGRP1 KO mice relative to WT (Figure 7A-B). We hypothesized that these cells may not be detected in RasGRP1 KO thymi because of earlier emigration of γδT17 TF^+^ cells into the periphery, but analysis of splenocytes and lymphocytes revealed this c-Maf^+^RORγt^+^Vγ2^+^ population trafficked to the LN in WT mice, and was not present in RasGRP1 KO mice in all tissues analyzed (Figure 7B). Furthermore, we identified that these immature c-Maf^+^RORγt^+^Vγ2^+^ cells were specifically trafficking to the inguinal LN over the mesenteric LN (Supplementary 2A).

**Figure 7:**
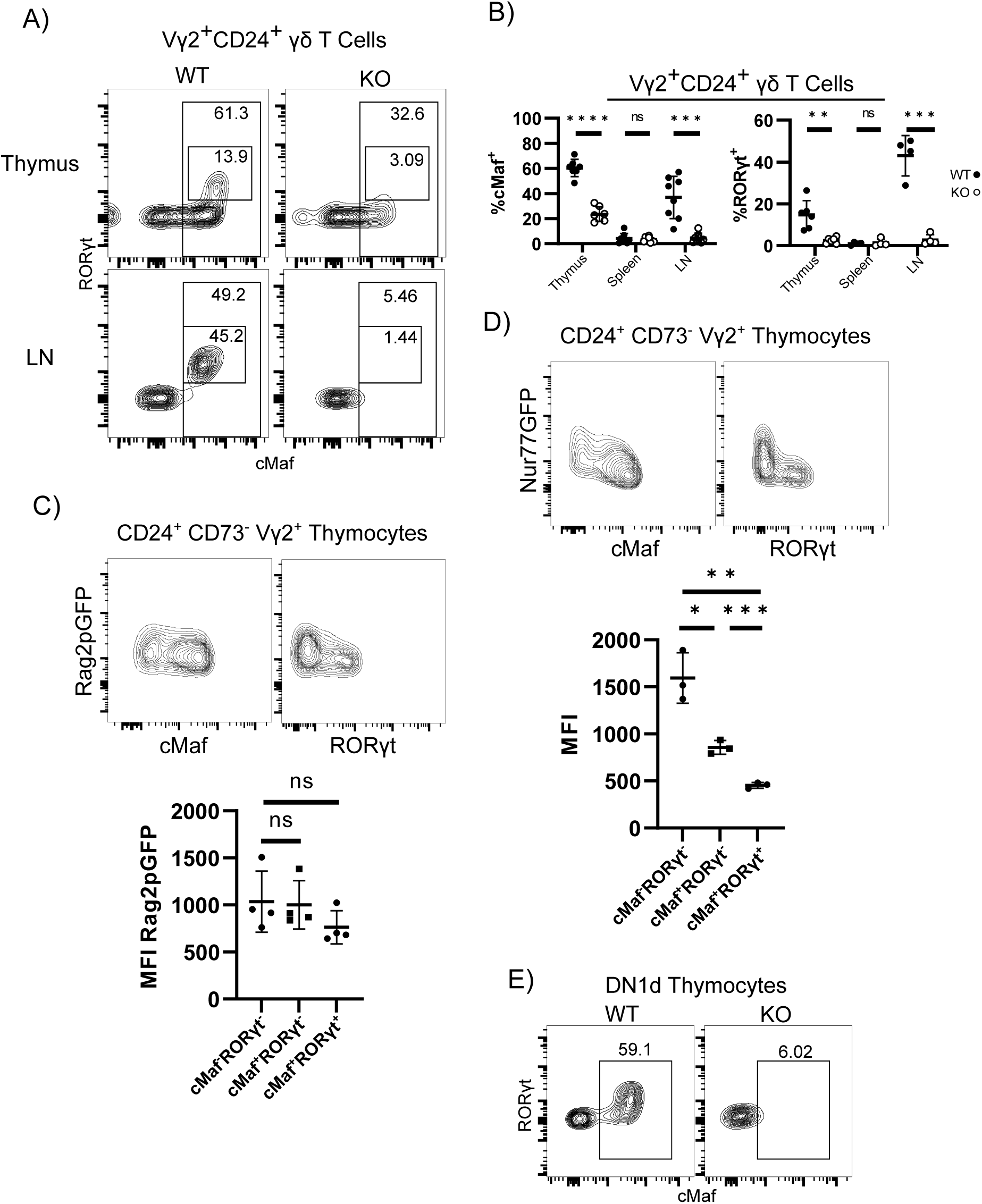
γδT17 programming defect is detectable among immature γδ thymocytes. A) FACS plots and B) dot plots showing c-Maf and RORγt expression in WT and RasGRP1 KO Vγ2+CD24+ thymocytes and circulating in the periphery. C-D) Flow plots (top) and dot plots (bottom) demonstrating the relationship between c-Maf, RORγt, and C) Rag2pGFP or D) expression in WT thymocytes. E) Representative FACS plots of c-Maf and RORγt expression in WT ot RasGRP1 KO DN1d thymocytes.All Nur77GFP mice were male. All data N = 3 or more, 3 or more independent experiments.

Our γδT17 TF analysis suggested that only Vγ2^+^ thymocytes that expressed c-Maf most strongly began to express RORγt (Figure 7A), and TCR signaling is thought to substantially direct γδ effector programming. Therefore, we were interested in examining the kinetics of c-Maf expression in these γδ thymocytes in terms of TCR expression and signaling. First we analyzed Rag2pGFP mice^16^ that express GFP under the control of the Rag2 promoter. In this model, GFP expression peaks during V(D)J recombination and intracellular GFP concentration decreases over time as a factor of protein turnover and proliferation. Analysis of Rag2pGFP mice found no significant difference in GFP intensity between immature c-Maf^+^ and c-Maf^-^ Vγ2^+^ thymocytes, but a non-significant trend towards immature c-Maf^+^RORγt^+^Vγ2^+^ thymocytes expressing GFP less intensely (Figure 7C). Next we analyzed Nur77^GFP^ mice, which express GFP as a factor of duration and intensity of TCR signals received^17^. Analysis of Nur77^GFP^ mice revealed a clear trend of less TCR signaling correlating with increased c-Maf expression, and immature Vγ2^+^ thymocytes that were RORγt^+^ expressed the lowest amount of Nur77^GFP^ (Figure 7D). Since our Nur77^GFP^ data suggested that c-Maf and RORγt expression correlated with low or no TCR signaling, and DN1d thymocytes have been reported to be pre-programmed to the γδT17 lineage in the absence of γδTCR signaling^23^, we extended our TF analysis to the DN1d population. Within the DN1d thymocyte population, RasGRP1 was critical for c-Maf expression even prior to TCR expression (Figure 7E). Taken together, these data suggest that RasGRP1 transmits crucial signals driving γδT17 programming in DN1d thymocytes prior to TCR expression and that programming continues in Vγ2^+^ thymocytes that receive low or no TCR signaling.

### CCR9 stimulation upregulates c-Maf in immature Vγ2+ thymocytes

Given that we observed c-Maf expression begins prior to TCR expression in DN1d thymocytes and only continues in cells receiving the lowest levels of TCR signals, we hypothesized that receptors other than the TCR may signal through RasGRP1 to induce c-Maf expression. Previous data from our lab demonstrated that RasGRP1 is required for effective activation of ERK and trafficking of thymocytes in response to some chemokines including CCL25^24^ and SDF1α^12^. Therefore, we analyzed immature γδ thymocytes for CCR9 expression, and induction of c-Maf following stimulation with CCL25. Immature CD73^-^ γδ thymocyte populations that expressed c-Maf also expressed the highest levels of CCR9, and RasGRP1 KO mice not only had lower c-Maf expression but also lower CCR9 expression as well (Figure 8A).

**Figure 8:**
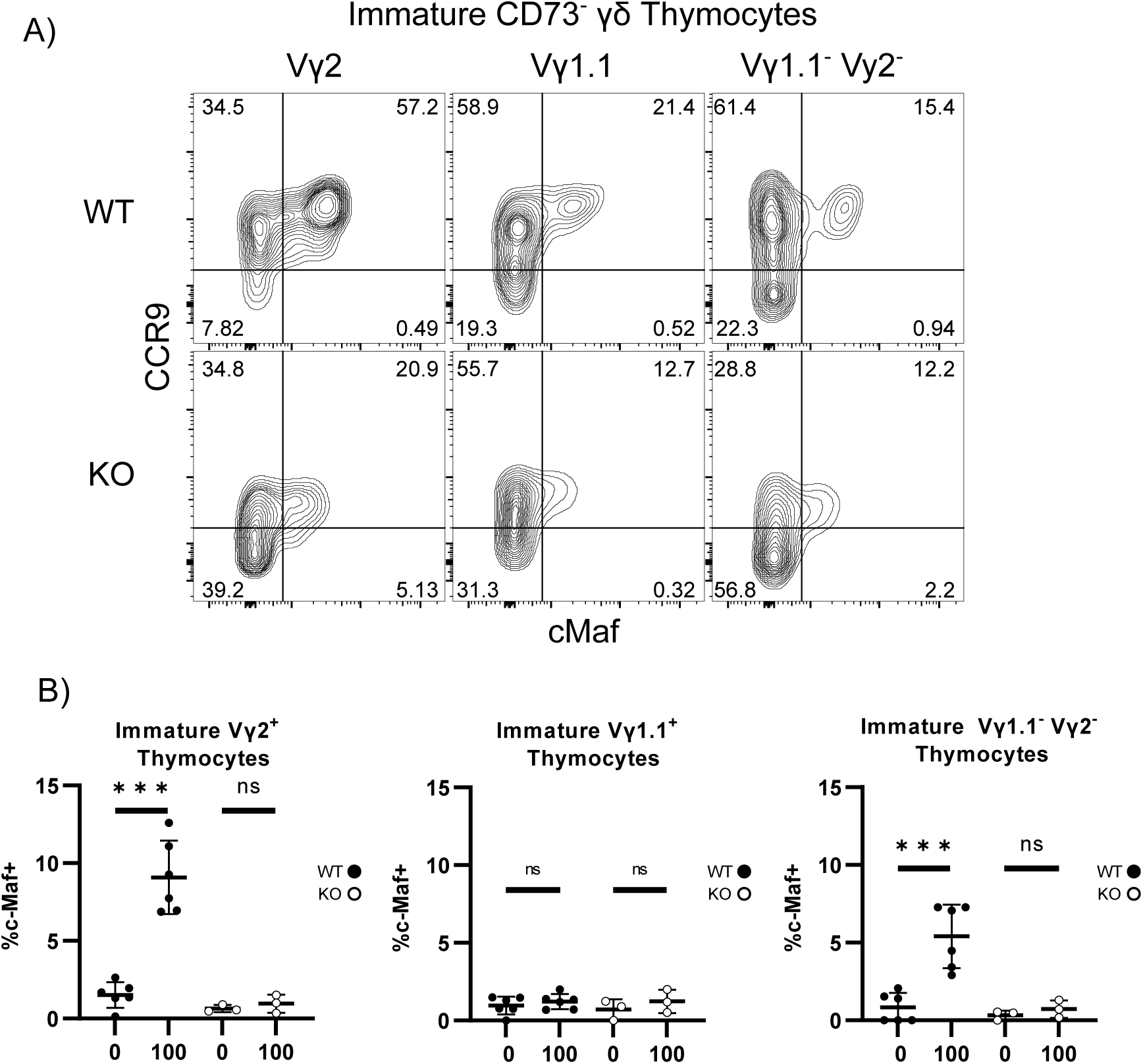
CCR9 Stimulation upregulates c-Maf in immature CD73- Vγ2+ thymocytes. A) Ex-vivo expression of CCR9 and c-Maf in immature γδ thymocytes. Data representative of N=4, 3 independent experiments. B) Expression of c-Maf in immature γδ thymocyte subsets after in-vitro culture for 1 day of rest followed by 1 day of medium or CCL25 stimulation at 100 ng/mL in WT or RasGRP1 KO Mice. WT N = 6, 5 independent experiments, KO N = 3, 3 independent experiments.

We next directly tested whether CCR9 stimulation could upregulate c-Maf in immature γδ thymocytes. Bulk thymocytes were incubated in complete media alone for 24 hours to allow c-Maf expression to decline to baseline then were subsequently stimulated with or without 100ng/mL of CCL25 in complete media for 24 hours. The addition of CCL25 caused a significant upregulation of c-Maf in both WT immature Vγ2^+^ and Vγ1.1^-^Vγ2^-^ γδ thymocytes, but not in the corresponding RasGRP1 KO γδ thymocytes (Figure 8B). These results suggest that CCR9 signaling through RasGRP1 contributes to the γδT17 programming of immature Vγ2^+^ thymocytes.

## Discussion

TCR signals received by T cells during development in the thymus are known to direct both the clonal selection and lineage commitment of αβ T cells, but a poor understanding of γδ T cell ligands has complicated research into how γδTCR signals, or signals from other receptors, contribute to γδ-T cell development. Generally, signals from the γδTCR are thought to direct development into the γδ-T cell lineage over the αβ lineage, and further γδTCR signals are thought to program the effector fate of γδ thymocytes. However, it remains unclear how signaling pathways downstream of the γδTCR and/or other receptors work together to translate cumulative signals into distinct effector lineages.

αβ T cells require RasGRP1 and SOS1 for efficient β-selection^12^, and evidence suggests that developing T cells that receive strong γδTCR signals instead of weaker pre-TCR signals are directed into the γδ lineage^3,25–27^. This lineage selection process has been found to be due to stronger signals through the γδTCR causing MAPK activation and upregulation of Id3^28^. The Ras pathway can be activated in developing T cells through either RasGRP1 or SOS1^9^, and efficient β-selection has been found to depend on both Ras GEFs^12,14^. Interestingly, we have found no evidence of diversion from the γδ lineage in RasGRP1 KO mice, which suggests that γδ thymocytes do not depend on RasGRP1 for their lineage instruction. This may be due to the γδTCR providing stronger signals than the pre-TCR that engage SOS1 signaling leading to commitment to the γδ-lineage. While RasGRP1 deficient mice have normal numbers of γδ thymocytes, SOS1 deficient mice have an ∼50% reduction in γδ thymocytes^10^. Importantly, whether SOS1 is involved in γδ-lineage commitment, proliferation, or effector programming during development is currently unclear.

Despite no change in numbers of developing γδ thymocytes in adult RasGRP1 KO mice, we did detect an expansion of circulating mature γδ T cells. This is consistent with mature γδ T cells in circulation proliferating in the lymphopenic RasGRP1 KO environment. While previous research has not suggested a proliferative increase in RasGRP1 KO mice^14^, our Ki67 analysis may be more sensitive because of segregating the analysis of γδ-T cells based on CD24 expression. Our WT/RasGRP1 KO mixed BMC experiments are consistent with this hypothesis in the periphery where RasGRP1 KO cells have no substantial advantage populating CD73^-^ γδ compartments. Mixed BMCs did have a strong WT advantage to populating CD73^+^ γδ compartments in the thymus and periphery, which may be due to either RasGRP1 KO thymocytes not receiving the required signals to enter the CD73^+^ lineage^6^, or failing to proliferate in response to stronger TCR signals during development^14^. Taken together, these data are consistent with RasGRP1 not having a substantial role in the bulk development or homeostatic proliferation of mature circulating γδ T cells.

Our lab and others^14^ have noted that RasGRP1 KO mice have an apparent deficiency in circulating CD44^+^CD62L^-^ γδ T cells. This population contains the same CD44^hi^ population that would be expected to contain γδT17s^7^. Therefore, we considered if the loss of Vγ2^+^ γδT17s in RasGRP1 KO mice was due to a failure to generate naive Vγ2^+^ γδs, or a failure to maintain the Vγ2^+^ γδT17 population once generated. Our experiments do not reveal any failure to generate naive γδ T cells in RasGRP1 KO mice, and in fact, the numerical expansion of that population is consistent with the general trend of an expanded mature γδ T cell population. Since ERK signals can function as pro-survival signals, we considered if the loss of T_EM_ γδ T cells was due to a failure to maintain that population once generated. However, a RasGRP1/Bim DKO condition did not rescue this T_EM_ population. Therefore, we conclude that RasGRP1 must be required to generate this population from either programming immature γδ thymocytes or polarizing naive γδ T cells.

Specific TCR signaling pathways have been implicated in programming γδ T cells for different effector fates^7,23,29^. In this study we have identified how signals passed through RasGRP1 contribute to the effector fate programming of γδT1s and γδT17s. Our findings concur with previous experiments that found an increase in γδT1s in the lymph nodes of RasGRP1 KO mice^14^, and we identified this expansion as early-life derived CD8^+^ γδT1s. These cells have been recently described by Sumaria and colleagues^20^, and were found to develop in greater numbers in the presence of a MEK1/2 inhibitor. Therefore, our findings suggest that in WT mice, RasGRP1 activates ERK to constrain CD8^+^ γδT1 development early in life.

γδT17s in mice have been thought to be produced primarily through a wave of programming predominately Vγ4^+^ and Vγ2^+^ thymocytes in the perinatal period^23,30^, and to an unknown extent through activation of naive Vγ2^+^ γδs in adults ^31,32^. Surprisingly, our data suggests that the two neonatal γδT17 populations have substantially different requirements for RasGRP1 for their development. While the Vγ2^+^ population of γδT17s is almost completely absent in RasGRP1 KO mice, a substantial Vγ1.1^-^2^-^3^-^ population that we assume to be Vγ4^+^ remains. This effect of RasGRP1 is similar to the Vγ2 specific loss of dermal γδT17s in SOX13 mutant mice^33^, which suggests a close mechanistic relationship between RasGRP1 signaling and SOX13 expression. In addition, a proportional loss of thymic resident γδT17s is consistent with a reduction in total Vγ4^+^ γδ thymocytes during the perinatal period. Those Vγ4^+^ γδ thymocytes that are present, which are presumably Vγ4^+^, seem able to be programmed into functional γδT17s in the absence of RasGRP1, albeit at a reduced efficiency. In addition, the total reduction in neonatal Vγ1.1^-^2^-^3^-^ thymocytes may suggest that this population relies on signals transduced by RasGRP1 at some point during early development for their genesis or proliferation.

Regarding adult-derived γδT17s, current published evidence has demonstrated that naive γδ T cells can be polarized into γδT17s *in-vitro* given TCR stimulation, IL-1β, and IL-23^32,34,35^, and *in-vivo* when vaccinated with TCR specific antigen^31^. However, Yang and colleagues have recently found evidence that thymic programming of Vγ2^+^ γδT17s continues into adulthood^36^, but adult-programmed γδT17s may not contribute to peripheral γδT17 populations. Our data demonstrates that this thymic γδT17 programming is dependent on RasGRP1, so if this process is the same as the fetal-derived γδT17 programming, this finding would explain the almost total loss of Vy2^+^ γδT17s in RasGRP1 KO mice. Furthermore, our data suggests that these adult-programmed γδT17s leave the thymus and preferentially traffic to the certain LN, including inguinal LN. It is unclear if this partial development process is simply the remnant of neonatal γδT17 development, or if some conditions can induce completion of this development process to produce the adult-derived tissue-resident Vγ2^+^ γδT17s that have been observed by others^37^.

Our experiments explain one way that RasGRP1 contributes to thymic γδT17 programming before TCR expression. Previous research has identified SOX13^+^ DN1d thymocytes as the precursors of γδT17s, and that DN1d thymocytes already express CCR9, c-Maf, and RORγt^23^. Here, we demonstrate that upregulation of these TFs is dependent on RasGRP1, however, the specific relationship between RasGRP1 and SOX13 remains undefined. We find that CCR9 stimulation can contribute to expression of the c-Maf, and our Rag2pGFP data suggest that RORγt expression temporally follows c-Maf expression, which is consistent with previous data suggesting that c-Maf expression precedes and licenses RORγt expression^8,38^. In addition, RasGRP1 KO thymocytes also appear to have reduced expression of CCR9, which could further contribute to reduced γδT17 programming.

Once the γδ TCR is expressed, our Nur77-GFP data suggests that TCR signals compete with γδT17 programming to further direct development. Our data demonstrates that Vγ2^+^ thymocytes that receive the lowest TCR signal are the ones that express c-Maf in the adult thymus, which is consistent with the theory that TCR signals bias γδ thymocyte development to γδT1 programming^6,7^. However, the remaining immature Vγ2^+^ thymocytes appear to receive higher TCR signaling, and presumably are the source of the naive-like c-Maf^-^Vγ2^+^ RTEs found in the periphery, but we cannot rule out the possibility that Nur77 expression in these cells is due to ERK activating signals from a source other than the TCR^39^. Therefore, our data indicate there may be a hierarchy of development in which increasing levels of TCR signaling produce γδT17s, Naive γδ T cells, then γδT1s.

In summary, our data demonstrates that RasGRP1 does not contribute substantially to instructing thymocytes into the γδ lineage over the αβ lineage, but plays some, perhaps proliferative, role in the proportion of Vγ2^+^ and Vγ4^+^ thymocytes. We show that RasGRP1 has roles in γδ effector programming as it constrains CD8^+^ γδT1 differentiation, and enhances γδT17 programming in Vγ2^+^ thymocytes early in life. Critically, the chemokine receptor CCR9 contributes to Vγ2^+^ γδT17 programming in a RasGRP1 dependent manner, preceding γδ-TCR expression, and establishes an environment for direction into the γδT17 lineage.

## Acknowledgements

The authors thank Dr. Robert Ingham for critical review of the manuscript. Flow cytometry experiments were performed at The University of Alberta Faculty of Medicine & Dentistry Flow Cytometry Facility, which receives financial support from the Faculty of Medicine & Dentistry and Canada Foundation for Innovation (CFI) awards to contributing investigators. Health Sciences Laboratory Animal Services, and Mr. Bing Zhang and Dr. Thaisa Sandini provided animal husbandry and technical assistance.

## Author contributions

K.J., D.P.G.,and T.A.B. designed the experiments. K.J, D.P.G., L.M.H.C. and T.A.B analyzed the data. K.J., D.P.G., L.M.H.C., R.G.K., and J.F.M performed the experiments. K.J., and T.A.B. wrote and edited the manuscript and D.P.G and J.F.M edited the manuscript.

