## Supplemental Figures for "RasGRP1 Signalling Programs Developing γδ-Thymocytes towards the γδT17 Lineage Through Control of c-Maf Expression"

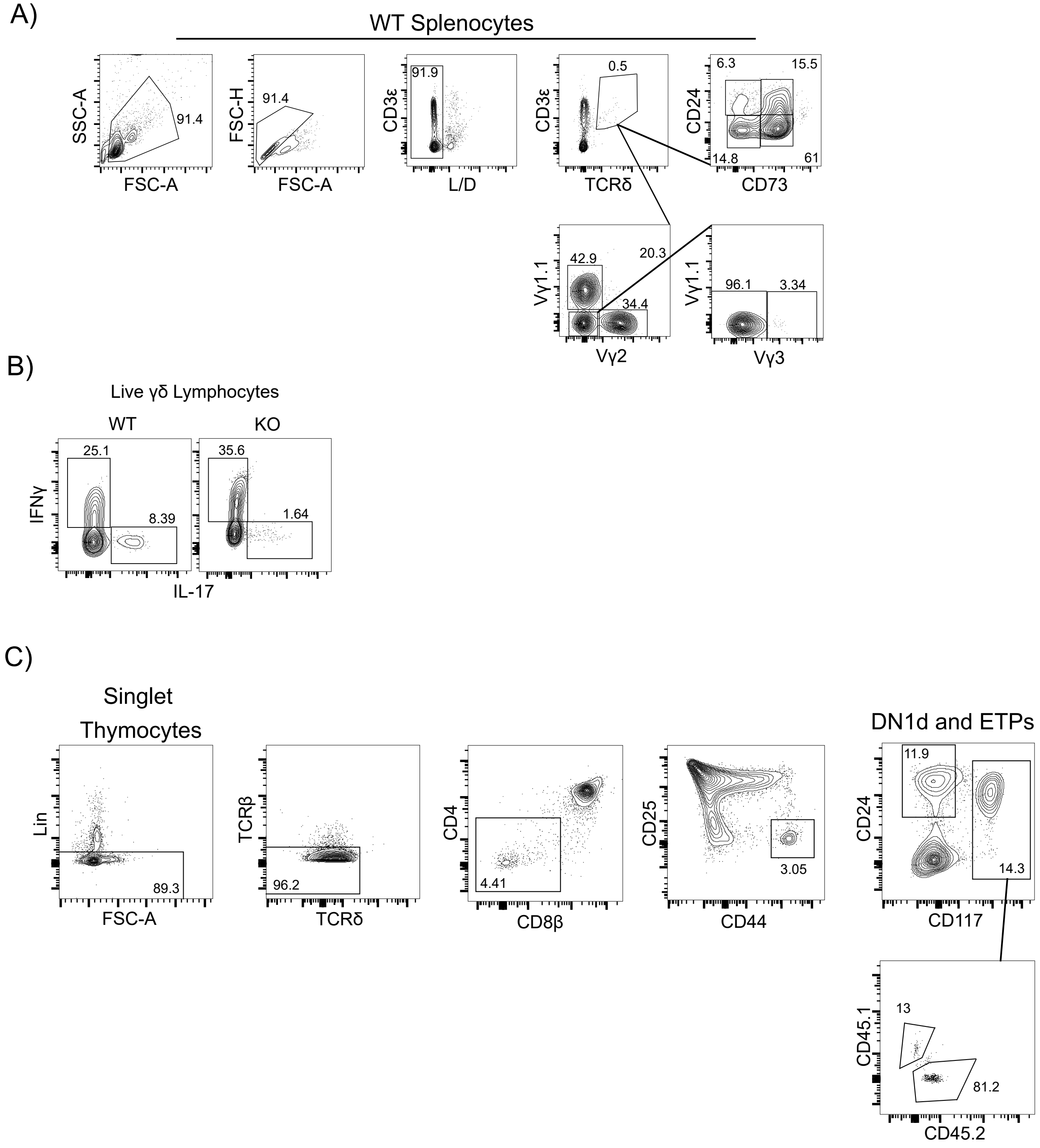


Supplementary Figure 1: Gating strategies for flow cytometry analysis. A) Gating for analysis of γδ T cells in terms of CD24/CD73, Vγ1.1/Vγ2/Vγ3. B) Gating of IL17 and IFNγ expression. C) Gating for ETP and DN1d thymocytes, and CD45 analysis of ETPs for BMCs.

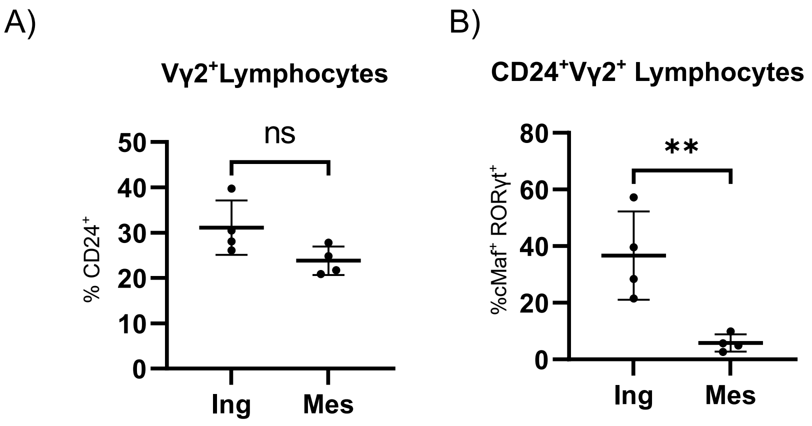


Supplementary 2: Immature c-Maf+RORγt+Vγ2+ lymphocytes bias trafficking towards specific LN. A) Dot plot of total Vγ2+ lymphocytes that are CD24+ in inguinal vs mesenteric LN. B) Dot plot of total CD24+Vγ2+ lymphocytes that are c-Maf+RORγt+ in inguinal or mesenteric LN. All data from 2 independent experiments, N=4.
